# Temporal Progression: A case study in Porcine Survivability through Hemostatic Nanoparticles

**DOI:** 10.1101/2021.05.25.445617

**Authors:** Chhaya Kulkarni, Nuzhat Maisha, Leasha J Schaub, Jacob Glaser, Erin Lavik, Vandana P. Janeja

## Abstract

This paper focuses on the analysis of time series representation of blood loss and cytokines in animals experiencing trauma to understand the temporal progression of factors affecting survivability of the animal. Trauma related grave injuries cause exsanguination and lead to death. 50% of deaths especially in the armed forces are due to trauma injuries. Restricting blood loss usually requires the presence of first responders, which is not feasible in certain cases. Hemostatic nanoparticles have been developed to tackle these kinds of situations to help achieve efficient blood coagulation. Hemostatic nanoparticles were administered into trauma induced porcine animals (pigs) to observe impact on the cytokine and blood loss experienced by them. In this paper we present temporal models to study the impact of the hemostatic nanoparticles and provide snapshots about the trend in cytokines and blood loss in the porcine data to study their progression over time. We utilized Piecewise Aggregate Approximation, Similarity based Merging and clustering to evaluate the impact of the different hemostatic nanoparticles administered. In some cases the fluctuations in the cytokines may be too small. So in addition we highlight situations where temporal modelling that produces a smoothed time series may not be useful as it may remove out the noise and miss the overall fluctuations resulting from the nanoparticles. Our results indicate certain nanoparticles stand out and lead to novel hypothesis formation.

## 1 Introduction

In this study, we use temporal analysis to discover patterns and generate insights from porcine data collected after the hemostatic nanoparticles are injected onto trauma-induced porcine animals. In particular we evaluate the state of the animal, in terms of survivability and blood clotting, over time as a result of the nanoparticle or control administration. Our study focused on the survivability factor along with clot formation components. 19 porcine animals were classified into three groups namely vehicle control, control nanoparticle and hemostatic nanoparticle groups. We evaluated this lab data as temporal models and then cluster the porcine animals based on their similarities and differences based on their reactions over time to the hemostatic nanoparticles and their survivability. Understanding when the animal moves from a trauma phase into a survivability phase or clotting phase is important to evaluate the impact of the nanoparticles. As a result, temporal analysis plays a key role here especially in understanding the overall trends in the animals based on the hemostatic nanoparticle or control administered. We present an exploratory study that used temporal analysis to find discrete intervals in time when the state of the animal transitions. However, we also outline situations where the modelling is useful and where it may not be as effective. We utilize this intuition to cluster the raw data to evaluate outcomes from this study in terms of animals exhibiting similar trends over time.

### Motivating Scenario

Traumatic injury is the leading cause of death in men, women, and children between the ages of 1 and 46 worldwide[1], and blood loss is the primary cause of death at acute time points post injury in both civilian[2] and battlefield trauma[3]. Immediate intervention is key to survival[4], and yet there are no reliable and readily available treatments for internal bleeding. We developed an intravenously infusible hemostatic nanoparticle that effectively stopped internal bleeding in a number of rodent models of trauma [5-7], but we have encountered off target effects in large animal (porcine) models of trauma that mimic infusion reactions seen in some people [8]. We developed a novel version to avoid these infusion reactions, and while it did not exhibit signs of complement activation seen previously, off target effects including non-specific clots were seen in some animals. To understand the features that correlated with these unexpected findings, we needed to be able to determine the specific features that correlated with the clots.

The porcine model is the gold standard for modeling trauma and critical for preclinical translation. It has been a model that is appreciated for how well it replicates the cardiovascular responses to trauma [9-11] and preclinical model for testing therapies [9-11, 22-27]. While the porcine model is exceptional for modeling cardiovascular responses, it has become a substantial question in the field as to whether they are appropriate for translation of nanomedicines because of the infusion responses [12, 13]. So, while one can obtain infusion responses to nanomedicines in mice and rats [18-20], the porcine response is a better predictor of the human response [14-17, 21].

As evident from the complexity of the reactions to hemostatic particles and variations possible by nanoparticles administered, it is imperative that we have a robust model to represent the changes to blood loss over time and identify similarities and dissimilarities between animals not just by treatment but also by the trends in their blood loss. In addition we also observe trends in the cytokine features that lead to reactions in the immune system that occur as a result of the trauma (including Il6, IL8, Neutrophils [8]). Understanding Cytokine reactions may also be useful in studying severe acute respiratory syndrome coronavirus [31,32]. There are several heterogeneous experiments in this framework that are conducted resulting in time series datasets with survivability or clotting outcomes. Understanding the time series representation for the progression of blood loss of the animal or cytokines helps understand the underlying complex mechanisms at play for the hemostatic nanoparticles administered to the animal. For example if a particle is administered to one animal and it behaves very similarly to other animals in a cluster in terms of its trend of survivability we can further explore what factors lead to the survivability potentially understanding complex cytokine mechanisms at play such as hemostasis (stopping of bleeding) or vasodilation (leading to more bleeding).

In this paper we present our preliminary work in representing the data in discretized temporal models [1]. We also utilize clustering to identify animals which are highly similar in these temporal models. We aim to produce novel hypotheses based on grouping animals based on their reaction to hemostatic nanoparticles.

Our overall contribution in this study is twofold in nature:

- We present multiple ways for bioinformaticians to be able to visualize the trends in blood loss as a result of administering the hemostatic nanoparticles.
- We have also shown that traditional smoothing techniques may sometimes hide the fluctuations in the data that are crucial in feature selection and understanding the mechanisms by which cytokine features can be impactful even in minor fluctuations.

We discuss our approach in section 2, followed by results in section 3.

## 2. Approach

The lab experiments involved intravenous administration of hemostatic nanoparticles in 19 porcine animals and their data was recorded. The blood loss data for each animal contained about 60 time instances, moreover each of these animals also had temporal data about the cytokine features.

These 19 animals were divided into 3 groups. The three groups consisted of control, hemostatic and vehicle control groups. Animals were observed after they underwent a trauma incision. Blood samples were collected and weighed at each minute to determine the amount of blood loss experienced by the animals. We generate temporal representations of the blood loss and cytokine features using exploratory models such as SMerg [28] and Piecewise Aggregate Approximation (PAA) [29]. Both SMerg and PAA generated representations of the blood loss experienced by the animals at every time interval that was helpful for the bioinformaticians to understand the overall trend in blood loss. PAA data proved beneficial in assessing which animals are similar within and across the different groups of control and vehicle control groups. Similarly, SMerg helped us see when the blood loss plateaued or dropped. SMerg provided a robust noise free representation of the data and we also clustered the outcomes from SMerg to observe which animals are in the same clusters. We performed a comparative analysis with variations of the clustering using SMerg and using the raw time series data.We found that the raw data captures some minor fluctuations which might be missed when the data is smoothed. Across all these comparisons we identified if animals consistently fell into similar clusters or fluctuated across methods in their cluster memberships. We also evaluated if there were a lot of cluster reassignments in certain nanoparticles which would indicate fluctuations in the data to warrant additional investigations in those nanoparticles in the lab.

Our exploratory data analysis is summarized in Figure 1 and the steps are discussed next:

**Figure 1:**
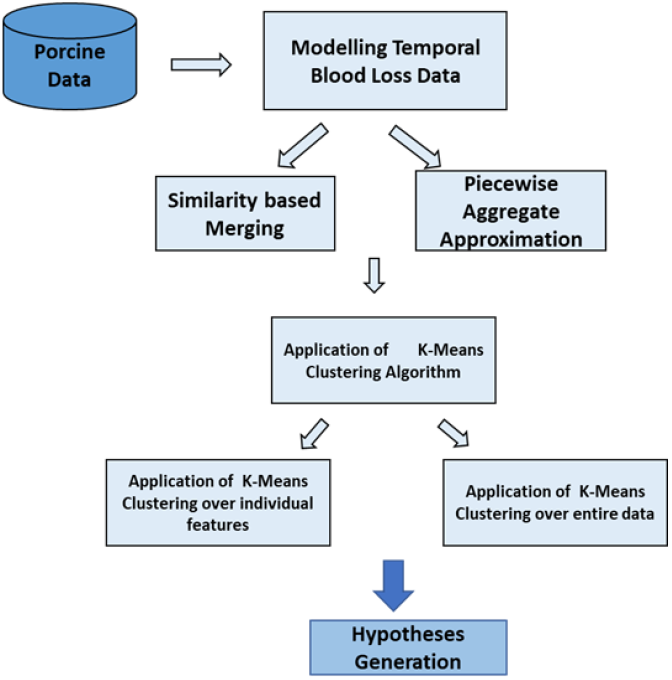
Evaluating Temporal progression.

### A) Similarity based Merging (SMerg)

In order to find representations in temporal blood loss data, we used our prior work on discovering temporal neighborhoods [28]. We have used this technique to divide the porcine data into temporal neighborhoods based on similarity in terms of blood loss experienced by animals at various time instances. Similarity based Merging (SMerg) uses a Markov model to generate a temporal neighborhood. It starts by dividing the temporal data into equal depth bins initially. These equal-depth bins are treated as the states of the Markov model. The distance between each bin and its neighboring bins is computed using distance metrics such as Mahalanobis, Kullback-Leibler distance measure (KL), and Bhattacharyya. Next, the Markov transition matrix is computed using a row-stochastic similarity matrix from the distance matrix. From the transition matrix, the adjoining bins with high degree of similarity are merged to generate the temporal neighborhood. This provides a set of discretized bins which represent the overall distribution of the data. the advantage of this method is that it smooths out the data without losing any key trends or patterns and provides a very robust summarization of the data

### B) Piecewise Aggregate Approximation (PAA)

PAA is a well-known dimensionality reduction technique in time-series mining [2]. It helps visualize the data in a condensed form. Input to the PAA algorithm is a time series. PAA divides the time series into a set of segments, and each segment is replaced by the mean of its data points. PAA proves to be helpful in identifying similarities and differences across the animals. PAA captures the essence of each time interval but shows it in a compressed manner. PAA replaces the time points in our data by the mean of those points. We have used PAA in our study to be able to monitor the movement of blood loss of porcine animals over continuous time intervals in a compact manner without losing sight of even a single rise and fall of data anywhere. PAA helps in recognizing anomalies in the data. SMerg on the other hand helps in generating plateaus and drops that help ascertain which nanoparticles elicit consistent behavior and till what point. Both PAA and SMerg are complementary in investigating the temporal progression of the survivability as a function of the blood loss in the porcine animals.

### C) K-Means Clustering

We conducted clustering experiments on the porcine data and created different types of inputs namely: SMerg Normalized data, SMerg Not normalized data with 2 values of k=5 and k=7 (based on the elbow method of determining an optimal K value with respect to Sum of Squared Errors). Here normalized data was created by comparing it to a baseline captured at the start of the experiments. We also considered raw data to evaluate whether SMerg is capturing the fluctuations that the bioinformaticians observed. We then computed the Sum of Squared Error values for blood loss and each cytokine measured feature namely IL6, IL8, Neutrophil. Our intuition was that the animals should fall into the same clusters regardless of the method used and if there are variations that would be interesting to evaluate further.

## 3. Experimental Results

### 3.1 Temporal Analysis

Temporal visualizations created using PAA and SMerg can be seen in figures 2 and 3 respectively. In figure 2 we can compare the original data seen in with the PAA (2(a)) and SMerg representations (2(b)) for Animal 4 that did not survive.. We can see that PAA helps in capturing the trend without losing all the fluctuations. Relative to PAA, SMerg is more suitable to smooth out the data to see overall plateaus and dips. SMerg smoothes out the data to capture the larger trends in the data. However, in this process it also leads to loss of minor fluctuations which might be interesting given the complexity of the reactions. Losing noise should generally not be an issue, however, if the data has minor fluctuations that need to be captured in that case a method like SMerg may not be useful.

**Figure 2:**
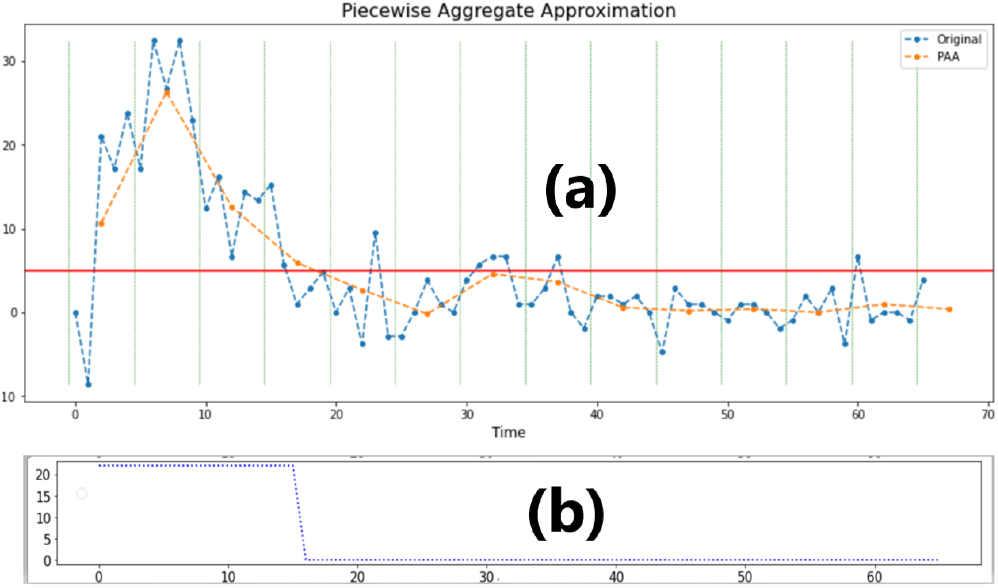
PAA and SMerg representation for Animal 4 of Group B (Not Survived)

**Figure 3:**
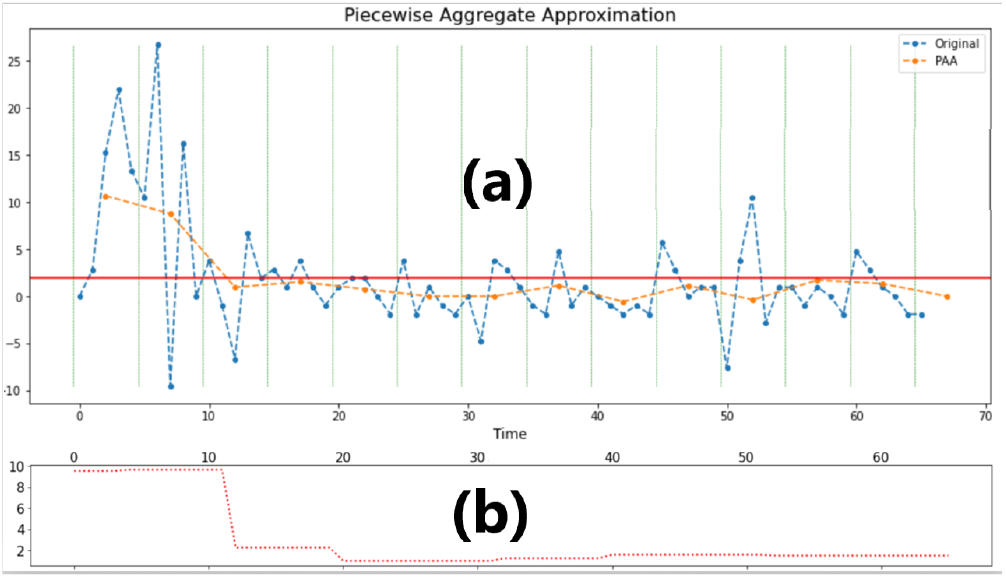
PAA and SMerg representation for Animal 6 of Group A (Survived)

As can be seen in figure 2, SMerg smoothes out the data. This smoothing is useful in seeing the dip and plateau when the Animal did not survive. However for the Animals that did survive the trends need to capture more nuance since fluctuations could have significance and show impact of the nanoparticles. In figure 3(b) SMerg analysis has a somewhat flat trend which may create a notion that the animal had a drop and plateaued (similar to figure 2 (b)). Figure 2 and 3 collectively help us understand that capturing noise is essential to get a comprehensive understanding of the data especially when used for decision making tasks such as predicting feature importance or associations.

### 3.2 Clustering

In figure 4 we depict SSE values for neutrophil, cytokines namely IL6, IL8 and blood loss using variations of data (after applying temporal modeling and raw data with or without normalization).

**Figure 4:**
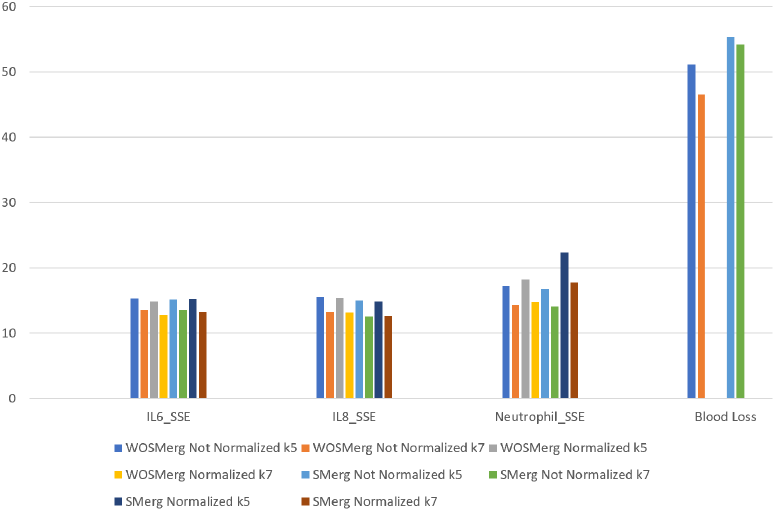
SSE across blood loss and Cytokine features.

We can see that blood loss has the highest SSE indicating the variance in the blood loss data. Neutrophil is the next highest SSE followed by IL6 and IL8.

We also capture how many times an animal switched clusters by normalization and by change in K values when we used the raw data. Figure 5 depicts each animal with the ‘number of changes’ measure. We can see that in most of the cases animals that have survived have more cluster switching as opposed to the animals that did not survive indicating that they do not have as much separation. This was quite interesting as this captured the minor fluctuations in the data as well. However, we did not see such cluster changing behavior in IL8 as shown in figure 6. In terms of cluster changes IL6 shows the most fluctuation however, in terms of SSE value Blood loss and Neutrophil exhibit a higher amount of fluctuation as shown in figure 7. We also see that IL6 and Neutrophils are ranked much higher in feature ranking as well, as shown in figure 8. Their activity in the trauma is critical to outcomes, and the cytokine levels.

**Figure 5:**
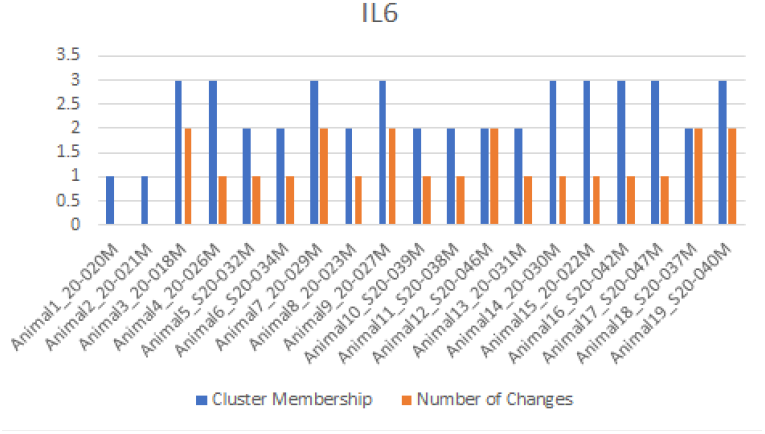
Cluster Membership for IL6 in raw data.

**Figure 6:**
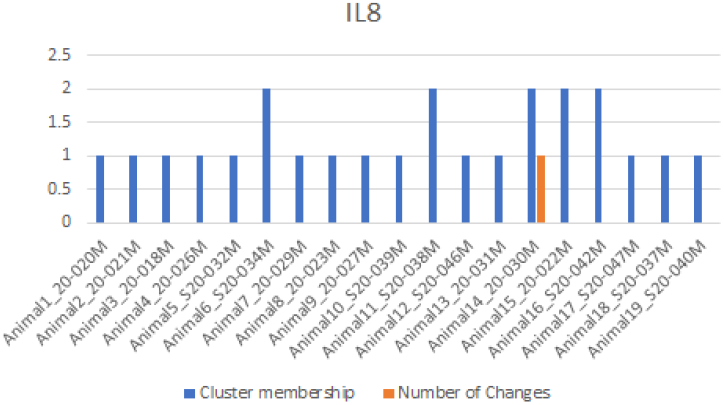
Cluster Membership for IL8 in raw data.

**Figure 7:**
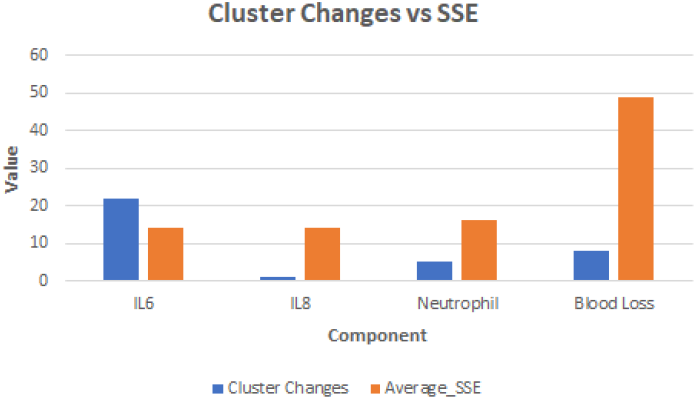
Cluster Changes vs SSE.

**Figure 8:**
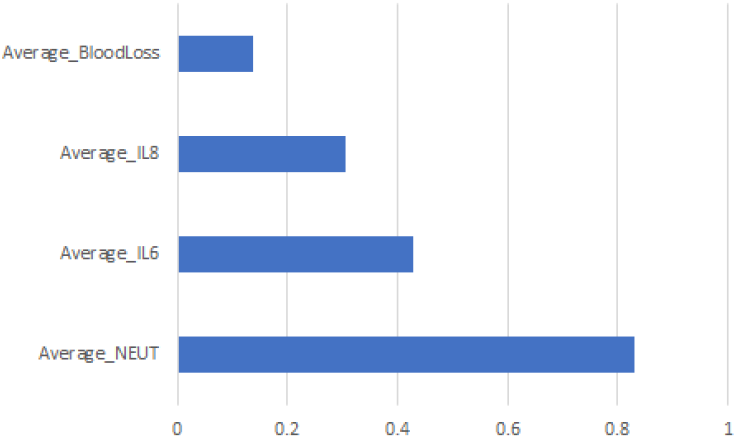
Feature Ranking.

## 4. Conclusions and Future Work

Visualizations created using PAA and SMerg were useful in making a general comparison among all the 19 animals in terms of temporal progression. We observed that if fluctuations are important to study underlying mechanisms, then the traditional time series methods may not be useful.Clustering captured deviations in IL6, IL8 and Neutrophils, which were also captured in feature raking. The cluster outcomes provided novel insights into these findings to observe why these animals survived and showed the fluctuations. We have explored supervised methods of using these findings to predict relationships to survival and clotting outcomes. The future work includes testing hypotheses based on the findings of IL6, IL8 and Neutrophils and their interactions in the lab.

## ACKNOWLEDGMENTS

Authors Lavik and Maisha would like to acknowledge AIMM Research award (DOD Award Number# W81XWH1820061).

